# Predicting post-TEVAR endoleaks: a pre-operative hemodynamic risk factor from patient-specific Fluid-Structure Interaction simulations

**DOI:** 10.64898/2026.03.16.712077

**Authors:** Francesca Duca, Silvia Tavarone, Maurizio Domanin, Daniele Bissacco, Santi Trimarchi, Christian Vergara, Francesco Migliavacca

**Affiliations:** LaBS, Department of Chemistry, Materials and Chemical Engineering ’Giulio Natta’, Politecnico di Milano, Piazza Leonardo Da Vinci 32, Milan, 20133, Italy; Department of Clinical Sciences and Community Health, Università degli Studi di Milano, Via Festa Del Perdono 7, Milan, 20133, Italy; Section of Vascular Surgery, Cardio Thoracic Vascular Department, Fondazione I.R.C.C.S. Ca’ Granda Ospedale Maggiore Policlinico, Via Francesco Sforza 35, Milan, 20133, Italy

**Keywords:** TEVAR, Endoleak, Thoracic aortic aneurysm, Fluid-Structure Interaction simulations

## Abstract

Thoracic Endovascular Aortic Repair (TEVAR) is a minimally invasive procedure for the treatment of thoracic aortic pathologies, such as Thoracic Aortic Aneurysm (TAA). Computational simulations can provide valuable insights into TEVAR outcomes and complications prior to surgery, making them a useful tool in the procedural planning. In this work, Fluid-Structure Interaction (FSI) computational simulations are carried out in ten pre-TEVAR patient-specific TAA cases, for which post-TEVAR outcomes are known, to quantify the hemodynamic drag forces acting on the aortic wall. Based on these results, this study proposes a new risk factor *R* to predict the occurrence of type I and III endoleaks. The patient cohort is divided in a calibration set, used to associate specific R values with three different risk levels, and a validation set, to test the risk factor efficacy. Based on the risk factor values obtained for the calibration set, *R* ***≤*** 0.33 is associated with low risk of endoleak formation, 0.33 ***<*** *R* ***≤*** 0.67 with moderate risk, and *R* ***>*** 0.67 with high risk. Once it is applied to the validation set,the risk factor is able to predict the formation of a type Ia endoleak. The risk factor proposed in this work is capable of identifying all the endoleak cases analysed, as well as conditions known to increase the risk of TEVAR complications. This study represents a preliminary attempt to determine whether pre-TEVAR hemodynamics can effectively predict post-TEVAR complications and thereby aid clinicians in the pre-operative planning.

## 1 Introduction

Since FDA approval in 2005 [1], Thoracic Endovascular Aortic Repair (TEVAR) has become a widely accepted and established alternative to open surgical repair for the treatment of a broad spectrum of thoracic aortic pathologies [2]. Specifically, TEVAR is a minimally invasive procedure in which a self-expanding stent-graft is introduced via a catheter and deployed in the Thoracic Aorta (TA), to exclude the diseased segment and restore physiological blood flow conditions [3]. To ensure long-term procedural efficacy, endovascular repair requires adequate stent-graft stability at both the proximal and distal device fixation sites, known as *Landing Zones* (LZs), with at least 2 cm of healthy aortic tissue [4].

Nevertheless, TEVAR outcomes are still limited by a significant risk of reintervention [5]. In the context of aneurysmal disease management, one of the main reasons for reintervention is the occurrence of *Endoleaks* (ELs), which are defined as persistent blood flow outside the stent-graft that continues to perfuse the aneurysmal sac [6]. In clinical practice, ELs are classified into five different types (for a more comprehensive overview of endoleaks, the reader is referred to Wanhainen et al. [7]). In this paper, we will limit our discussion to two specific types: *type I ELs*, which occur because of an ineffective seal between the proximal (*IA*) or distal (*IB*) LZ and the vessel wall, and *type III ELs*, which are caused by a structural failure of the endograft itself, usually due to the separation of the modular parts of a multi-component stent-graft [8]. Post-TEVAR ELs have been proven to be associated with large aortic diameter and tortuosity, suboptimal LZ length, proximal landing in 0-2 zone, Left-Subclavian Artery (LSA) coverage, large stent-graft diameters, excessive oversizing, bird-beak configuration, and the number of stent-grafts deployed [9–13].

Computational simulations can provide valuable insights into TEVAR outcomes and complications prior to surgery, making them a useful tool in the procedural planning [14]. Structural analysis has been widely used to simulate stent-graft deployment in several clinical applications [15–19], including TA pathologies such as penetrating aortic ulceration [20], dissection [21], and Thoracic Aortic Aneurysm (TAA) [22, 23]. On the other hand, Computational Fluid Dynamics (CFD) allows for the quantification of the hemodynamic *Drag Forces* acting on both the aortic wall and the endograft [24–27]. Previous studies have shown that both drag forces magnitude and orientation can help identify biomechanically hostile LZs, which may lead to poor stent-graft sealing or migration [28, 29]. Figueroa et al. [30] combined CFD and structural modeling to assess the drag forces acting on the stent-graft, demonstrating that computational methods can enhance the understanding of the loads experienced *in vivo* by the endograft, and therefore improve TEVAR outcomes. Prasad et al. [31] integrated CFD and structural analysis to study the drag forces on a multi-component stent-graft in a patient-specific TAA case, finding that opposing forces at the interface of two modular parts were linked to a type III EL occurring four years after the procedure. Notwithstanding, since many relevant mechanisms involve interaction between blood flow and the aortic wall, Fluid-Structure Interaction (FSI) modeling may offer a more accurate representation. Despite its potential, FSI remains underused in TEVAR studies due to its high computational cost [32–34], and to date, no studies have computed the drag forces using FSI to explore their association with TEVAR outcomes.

In this work, we apply the FSI computational framework presented in [35] to ten patient-specific TAA cases, for which post-TEVAR outcomes are known through follow-up Computer Tomography Angiography (angio-CT) scans. This study aims to:(i) quantify the drag forces acting on the aortic wall prior to the intervention and, based on these results, to develop a new hemodynamic risk factor that may predict the formation of post-TEVAR type I and III ELs; (ii) validate the predictive capability of the risk factor by comparing its predictions with the post-TEVAR follow-up outcomes (i.e., presence or absence of type I and III ELs) of each patient. To this end, the patient cohort is divided into two groups: an eight-patient calibration set, used to associate specific values of the risk factor with three levels of risk (i.e., low, moderate and high), and a two-patient validation set, employed to validate blindly the predictive capability of the risk factor. The ultimate goal is to determine whether pre-TEVAR hemodynamics can effectively predict post-TEVAR complications and thereby aid clinicians in the pre-operative planning.

## 2 Methods

From this point onward, we refer to: the *Proximal Landing Zone* as PLZ, i.e., the portion of healthy TA where the stent-graft proximal end is anchored to the aortic wall; the *Distal Landing Zone* as DLZ, i.e., the portion of healthy TA where the stent-graft distal end is anchored to the aortic wall; and the *Overlap Landing Zone* as OLZ,i.e., the region where two modular parts of a multi-component stent-graft overlap with each other.

### 2.1 Clinical data

A cohort of ten patients, who underwent TEVAR for a diagnosis of TAA without mention of rupture, was selected for this study. All patients were treated at the Section of Vascular Surgery of Fondazione I.R.C.C.S. Ca’ Granda Ospedale Maggiore Policlinico in Milan. For all patients, pre-operative, post-operative, and follow-up angio-CT scans were available, with a mean follow-up duration of 1.5 years (*±*1.3 years). Angio-CT images acquisition was performed with a Siemens Somatom Definition Flash scanner (Siemens Healthineers, Forchheim, Germany) with the following acquisition parameters: slice thickness of 3.0 mm, reconstruction matrix of 512 × 512 pixels and in-plane resolution of 0.629 mm × 0.629 mm. All patients’ data were anonymized by a staff physician before the computational analysis, and all patients provided informed consent for the treatment and the use of anonymized clinical data.

At the time of the last follow-up, five patients (i.e., PT-09, PT-22, PT-23, PT-40 and PT-78) showed no complications related to the TEVAR procedure, whereas the remaining five (PT-10, PT-21, PT-50, PT-51 and PT-67) presented with ELs. More specifically, PT-21 developed a type IA EL, PT-50 presented a type III EL, PT-67 had a type IB EL, PT-10 exhibited both type IB and III ELs, and finally PT-51 showed both type IA and III ELs. The clinical characteristics of each patient under study are detailed in Table 1, while Figure 1a showcases the patient cohort, highlighting each PLZ, DLZ and OLZ, together with the location of the ELs.

**Table 1:**
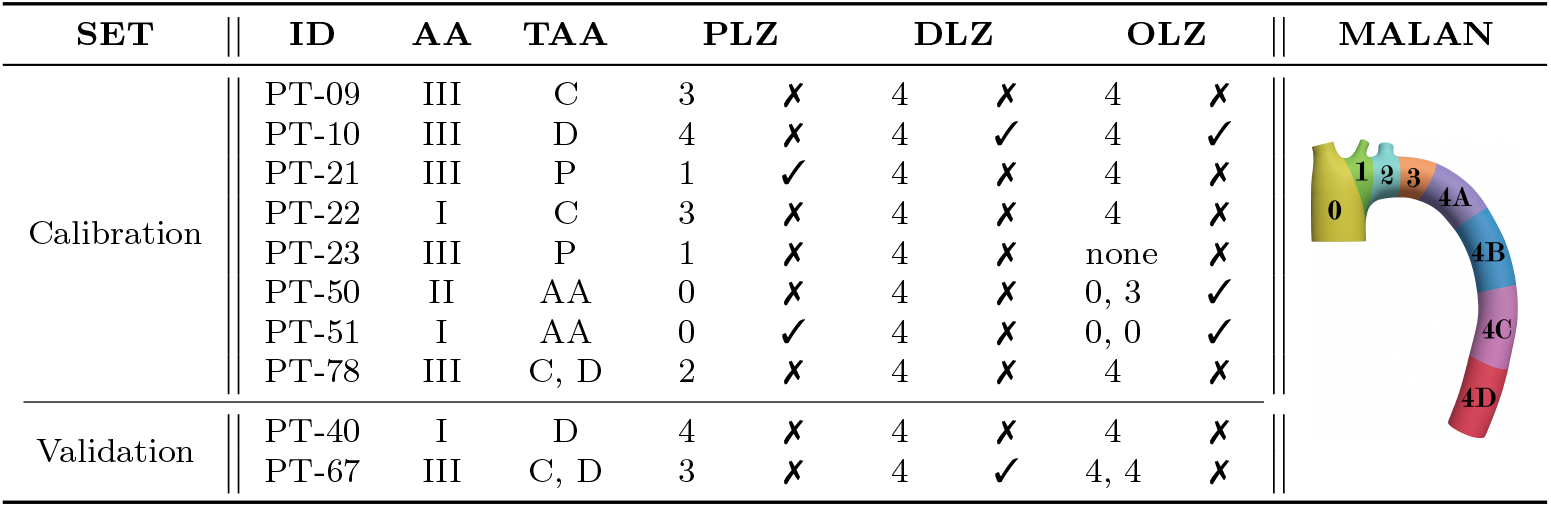
Patient clinical characteristics. ID: Patient ID; AA: aortic arch type according to the Aortic Arch classification [36]; TAA: thoracic aortic aneurysm location along the descending thoracic aorta (Proximal (P), Central (C), Distal (D), or Aortic Arch (AA)); PLZ, DLZ, OLZ: proximal, distal and overlap landing zones chosen by the clinicians during TEVAR. Left semi-column under PLZ, DLZ and OLZ: number of the landing zone where the corresponding sealing is performed, according to the MALAN classification [37] (see the last column of the table). PT-23 has only one stent-graft, whereas PT-50, PT-51 and PT-67 present three stent-grafts and therefore have two OLZs. Right semi-column under PLZ, DLZ and OLZ: post-TEVAR follow-up outcomes, i.e., presence (✓) or absence (✗) of endoleaks.

**Fig. 1:**
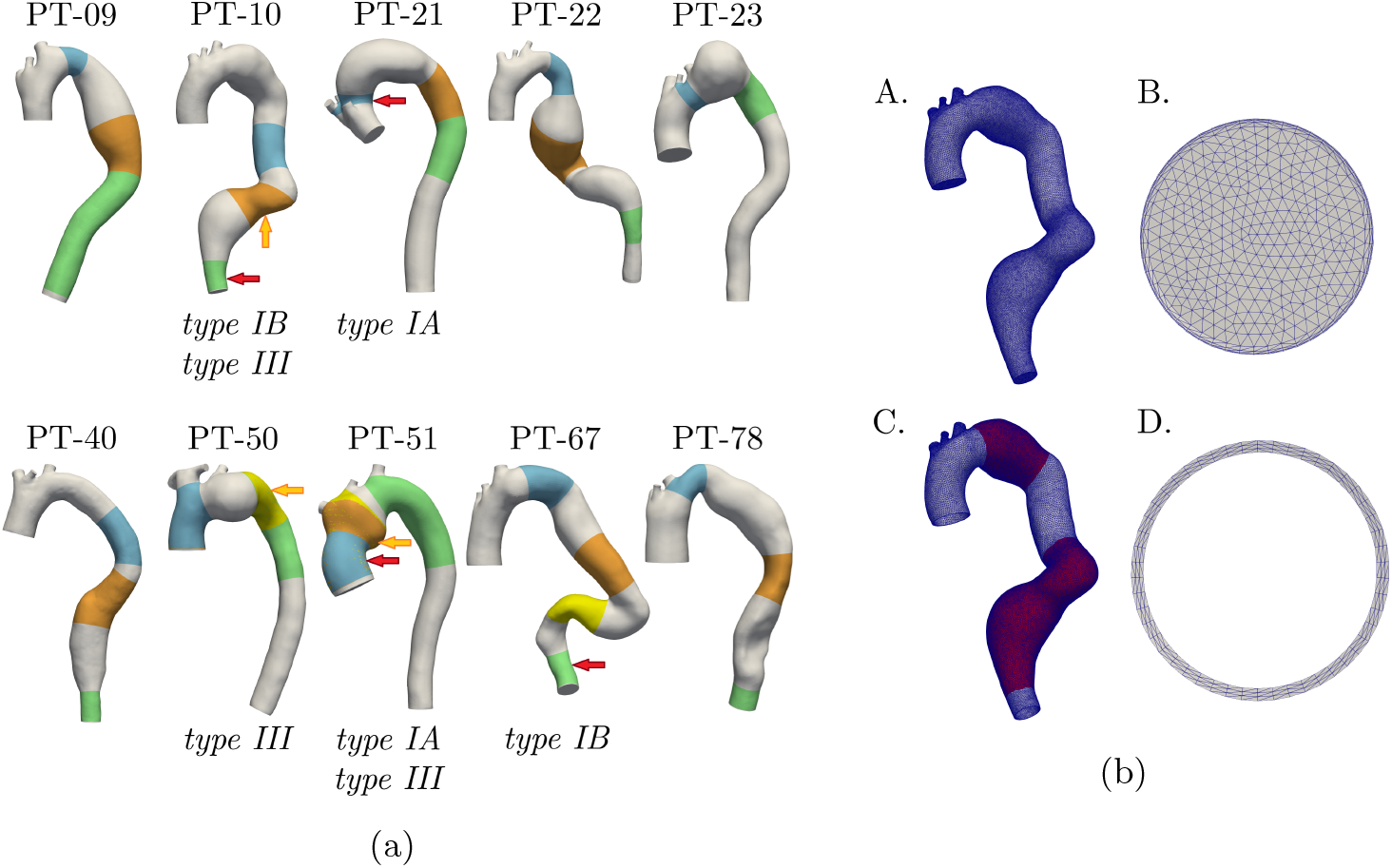
(a) Pre-TEVAR patient-specific geometries. Red arrows indicate the location of type I endoleak, while yellow arrows indicate the location of type III endoleaks. Proximal landing zones are highlighted in blu, distal landing zones in green, and overlap landing zones in orange and yellow. (b) Computational meshes of PT-10 (chosen here as representative). A. Fluid mesh. B. Zoomed-in view of the three-layered boundary layer. C. Structural mesh. D. Zoomed-in view of the three-layered structural mesh.

### 2.2 Computational meshes generation

Each computational domain is generated according to the pipeline described in Duca et al. [35]. For the sake of conciseness, only a brief summary is provided here.

First, the TA lumen of each patient is reconstructed from the pre-TEVAR angio-CT scans using a level-set segmentation technique with colliding fronts initialization, as implemented in the Vascular Modeling Toolkit (VMTK, http://www.vmtk.org [38]). The segmentation is performed from the Ascending Thoracic Aorta (ATA) to the origin of the coeliac tripod (see Figure 1a).

Subsequently, a volumetric fluid mesh is generated using VMTK (Figure 1b, panel A). For all patients, tetrahedral fluid meshes are built with a radius-dependent element size: the grid element size decreases as the diameter of the vessel diminishes, except for the aneurysm, where a constant value of 1.5 mm is imposed. Additionally, a boundary layer is generated, consisting of three sub-layers with a total thickness of 0.5 mm for the supraortic vessels and 1.0 mm for the rest of the domain (Figure 1b, panel B).

The structural meshes (Figure 1b, panel C) are constructed by extruding outward the fluid mesh’s external surface. The wall thickness is set to 10% of the lumen radius for the healthy vessel region and to a fixed value of 1.5 mm [39] for the aneurysmatic wall. For each patient, a three-layered solid mesh is generated (Figure 1b, panel D).

For both the fluid and the solid mesh, the mesh size is chosen after a mesh convergence analysis previously performed in Duca et al. [35].

### 2.3 Mathematical and numerical methods

All mathematical and numerical methods adopted in this study have been described in detail in our previous work [35]. Here, only the most relevant aspects are briefly summarized.

Blood is assumed to be a Newtonian, homogeneous, and incompressible fluid, with density *ρ*_f_ = 1060 kg m^*−*3^ and viscosity *µ*_f_ = 3.5 × 10^*−*3^ Pa s. Its behaviour is mathematically modelled with the Navier-Stokes equations written in the Arbitrary Lagrangian-Eulerian (ALE) formulation, where finite elasticity is used as lifting operator to recover the fluid domain configuration. To account for transition to turbulence, particularly relevant in presence of TAA [40], the *σ*-model Large Eddy Simulation (LES) turbulence model is adopted [41, 42].

The aortic wall is assumed to be nearly incompressible, with density *ρ*_s_ = 1000 kg m^*−*3^ and Poisson’s ratio *ν*_*s*_ = 0.45, and its dynamics is modelled using linear elasticity. The Young’s modulus of the healthy portion of the TA is set equal to 0.8 MPa [43],while for the aneurysmal region, a higher value of 1.2 MPa is imposed [44, 45] (see the region highlighted in red in Figure 1b, panel C).

The Boundary Conditions (BCs) applied to the FSI model are described as follows (Figure 2). For the fluid problem, a physiological time-dependent pressure wave with a period of 0.735 s and ranging between 70 and 120 mmHg is imposed at the ATA inlet (Figure 2b). At each supraortic outlets, namely the Brachiocephalic Artery (BCA), Left Common Carotid Artery (LCCA) and Left Subclavian Artery (LSA), an absorbing boundary condition consisting of a single resistance is prescribed (i.e., R_BCA_, R_LCCA_ and R_LSA_) (Figure 2c). Finally, at Descending Thoracic Aorta (DTA) outlet, a 3-element Windkessel model, composed by two resistances R_1_ and R_2_ and a compliance C, is applied (Figure 2a on the left). A non-zero outflow pressure P_out_ is set equal to 70 mmHg [46] (Figure 2a on the left). For the structural problem, at each inlet and outlet, a null displacement condition is prescribed, whereas on the external surface, a Robin condition, with springs of stiffness K_ext_ and elements of capacity C_ext_, is applied to model the constraint exerted by the surrounding tissue on the vessel’s movements (Figure 2a on the right). For the setting of the BCs parameters, we follow the criteria proposed in Duca et al. [35]. In particular, the supraortics resistances are individually calibrated for each patient, while the parameters R_1_, R_2_, C, K_ext_ and C_ext_ are kept constant across all patients and set equal to: R_1_ = 2.30 × 10^7^ Pa s m^*−*3^, R_2_ = 6.08 × 10^7^ Pa s m^*−*3^, C = 8.00 × 10^*−*7^ m^3^ Pa^*−*1^, K_ext_ = 1.50 × 10^6^ Pa m^*−*1^ and C_ext_ = 4.00 × 10^5^ Pa s m^*−*1^. Table 2 shows the final calibrated values of R_BCA_, R_LCCA_ and R_LSA_ for each patient.

**Fig. 2:**
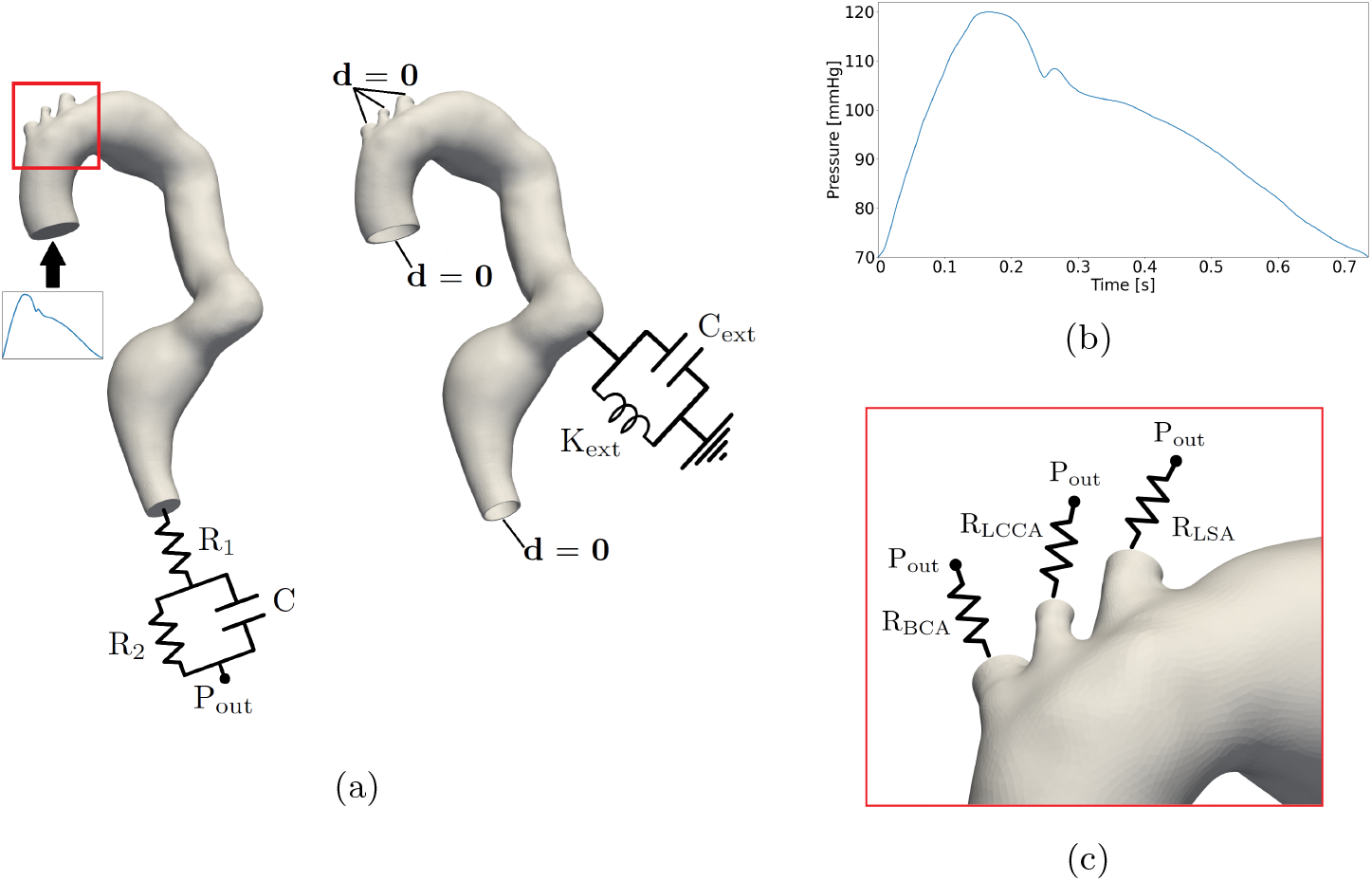
(a) Boundary conditions for the fluid (left) and the structural (right) sub-problems. (b) Physiological pressure wave imposed as inlet boundary condition. (c) Zoomed-in view of the boundary conditions imposed at the supraortic outlets.

**Table 2:** Boundary conditions parameters imposed at the supraortic outlets for all the patients under study.

| ID | $\mathbf{R}_{BCA}$<br>($\times 10^8 \text{ Pa s m}^{-3}$) | $\mathbf{R}_{LCCA}$<br>($\times 10^8 \text{ Pa s m}^{-3}$) | $\mathbf{R}_{LSA}$<br>($\times 10^8 \text{ Pa s m}^{-3}$) |
| --- | --- | --- | --- |
| PT-09 | 1.50 | 7.50 | 2.50 |
| PT-10 | 1.00 | 4.00 | 3.00 |
| PT-21 | 2.00 | 2.20 | 3.50 |
| PT-22 | 1.50 | 4.80 | 2.50 |
| PT-23 | 2.00 | 4.00 | 2.50 |
| PT-40 | 1.50 | 6.50 | 2.50 |
| PT-50 | 1.25 | 4.25 | 3.50 |
| PT-51 | 1.60 | 4.20 | 1.80 |
| PT-67 | 2.50 | 4.50 | 1.50 |
| PT-78 | 1.25 | 4.25 | 3.25 |

For the time discretization of the FSI continuous problem, a time-step Δt = 5 × 10^*−*4^ s is prescribed. A semi-implicit second-order Backward Differentiation Formula (BDF) is employed for the fluid sub-problem, with an explicit treatment for the turbulent viscosity. For the structural sub-problem, a first-order explicit BDF scheme is used. At each time step t^*n*+1^, the lifting problem is first solved, using the boundary displacement obtained from the structural solution at time t^*n*^, in order to update the fluid domain. Subsequently, the FSI problem is solved monolithically using GMRES, preconditioned via a block preconditioner.

For all patients, simulations are run over ten cardiac cycles, with the first three cycles discarded to allow the stabilization of flow velocity and pressure fields. All the numerical simulations are performed using life^x^ [47, 48]: a high-performance C++ library for the finite element simulations of multi-physics, multi-scale, and multi-domain problems, developed at the MOX, Department of Mathematics, with the collaboration of LaBS, Department of Chemistry, Materials and Chemical Engineering (both at Politecnico di Milano). Simulations are run in parallel on 112 cores of Intel Xeon Platinum 8480+@2.0 GHz CPUs, using the DCGP partition of the LEONARDO supercomputer (https://www.hpc.cineca.it/systems/hardware/leonardo/), available at the CINECA high-performance computing center (Italy).

### 2.4 Novel hemodynamic risk factor

The novelty of this study is the design of a pre-operative hemodynamics risk factor which may be able to predict type I and III ELs formation after TEVAR. This risk factor is based on the drag forces exerted by the blood flow on the aortic wall, computed at the systolic peak for each LZ surface area A as follows:

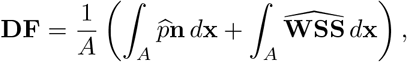

where 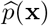 is the ensemble^1^ pressure at the systolic peak, 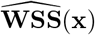 is the systolic ensemble wall shear stress vector, and **n** is the outward normal evaluated at the wall [35].

For the development of the risk factor, we hypothesize that, once the stent-graft is implanted, a high **DF** perpendicular component *DF*_⊥_ (see Figure 3a) could press the stent-graft against one side of the vessel wall, potentially detaching it from the opposite wall and thereby causing a type I EL. Moreover, this condition might lead to excessive vessel dilatation, resulting in a device overexpansion, causing a reduction in its radial force, and thus possibly compromising the stent-graft sealing. Finally, it has been proven that great *DF*_⊥_, caused by pronounced curvature or tortuosity, may induce lateral movements of the stent-graft, potentially leading to adverse events, such as type III EL [49]. On the other hand, a high **DF** parallel component *DF*_∥_ (see Figure 3a) indicates that the stent-graft could be subjected to a significant force along the direction of the blood flow, thus promoting device migration. This phenomenon could be particularly relevant in cases of type III ELs, as it may lead to downstream migration of the most distal stent-graft.

**Fig. 3:**
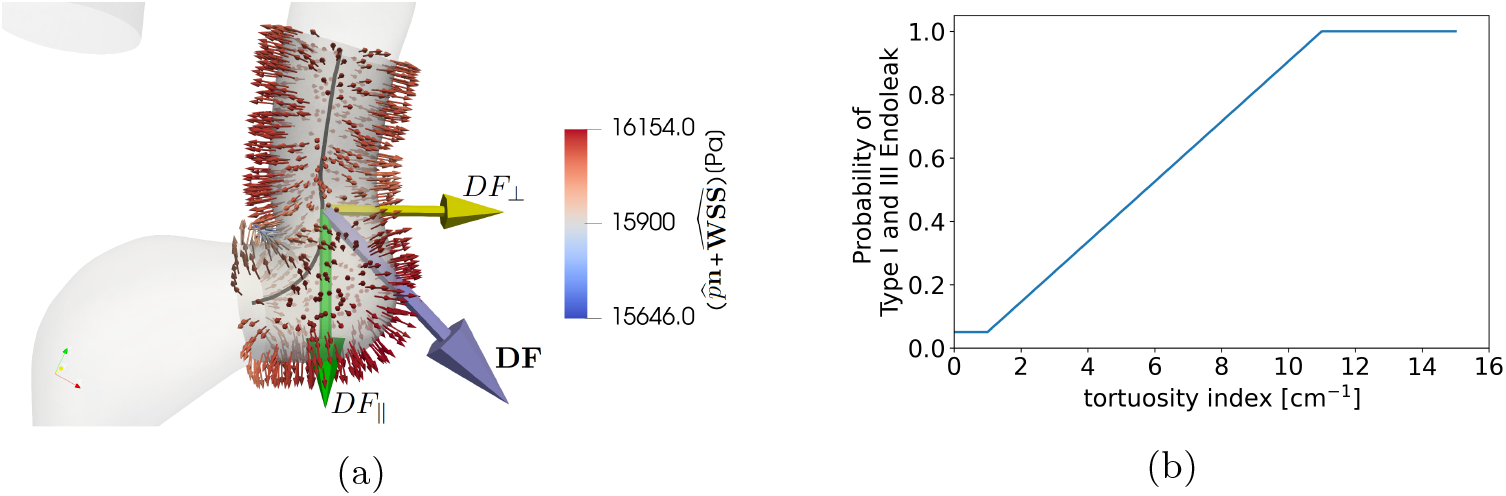
(a) Example of computation of the drag force components *DF*_∥_ and *DF*_⊥_ in PT-10 (shown here as a representative example). The gray line represents the center-line of the landing zone. Lilac arrow: drag force normalized by the landing zone area [N m^*−*2^]. Yellow arrow: normalized drag force perpendicular component *DF*_⊥_ [N m^*−*2^].Green arrow: normalized drag force parallel component *DF*_∥_ [N m^*−*2^]. (b) Probability of type I and III endoleak formation as a function of the tortuosity index of a certain landing zone, computed following Equation (1).

Another important aspect to consider is the TA tortuosity, as higher aortic tortuosity has been associated with higher incidence of both type I and III ELs [50]. For this reason, the risk factor proposed in this work should include also the probability *p*_*tortuosity*_ of type I and III EL occurrence as a function of the LZ tortuosity. The latter was statistically quantified by Ueda et al. [51], who conducted a retrospective study to analyse the correlation between tortuosity and EL formation. Using their published results as a reference, we define (see Figure 3b):

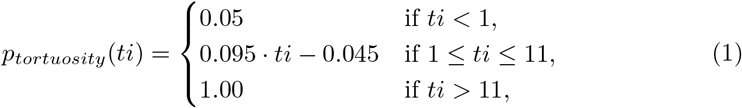

where *ti* is the tortuosity index; the latter is calculated as the sum of curvature values greater than 0.2 cm^*−*1^ measured every millimeter along the centerline in the LZ of interest [51, 52].

To build the risk factor *R*, we proceed as follows:

i. for each patient, the actual PLZ, DLZ and OLZ of the implanted stent-grafts are identified on the post-TEVAR angio-CT scans.
ii. A local risk factor *r* is computed for an initial 2-cm segment of each actual LZs, using the formula:

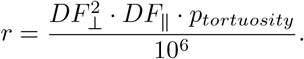 Here, *DF*_⊥_ is squared, as we hypothesize that it influences the occurrence of both type I and III ELs, whereas *DF*_∥_ primarily contributes to type III EL formation. For the PLZ, the initial segment is defined starting from the proximal aneurysm neck and extending upstreamly; for the DLZ it is defined from the distal neck and extending downstreamly; for the OLZ, it starts from the distal end of the most proximal stent-graft and extending upstreamly. The initial length of 2 cm is selected in accordance with current guidelines, which indicate this value as the minimum required for a secure stent-graft fixation [7].
iii. The initial segment is progressively extended by adding, along a local coordinate *z* (see Figure 4b and 4c), contiguous 0.5-cm sub-segments until the full extent of the actual LZ is reached. In cases where the actual LZ is shorter than 4 cm, a total length of 4 cm is nonetheless analysed, unless the end of the geometry is reached. Each sub-segment is created by cutting the aortic geometry with a plane perpendicular to the vessel centerline. *r* is computed for each incremental segment of each LZ.
iv. The risk factor *R* is computed by scaling each *r* value of each segment to the interval [0, 1], with the formula:

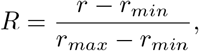

where *r*_*min*_ and *r*_*max*_ are the minimum and the maximum *r* values among all segments, respectively. Since PLZ, DLZ and OLZ are treated separately, *r*_*min*_ and *r*_*max*_ are identified independently for each case.

**Fig. 4:**
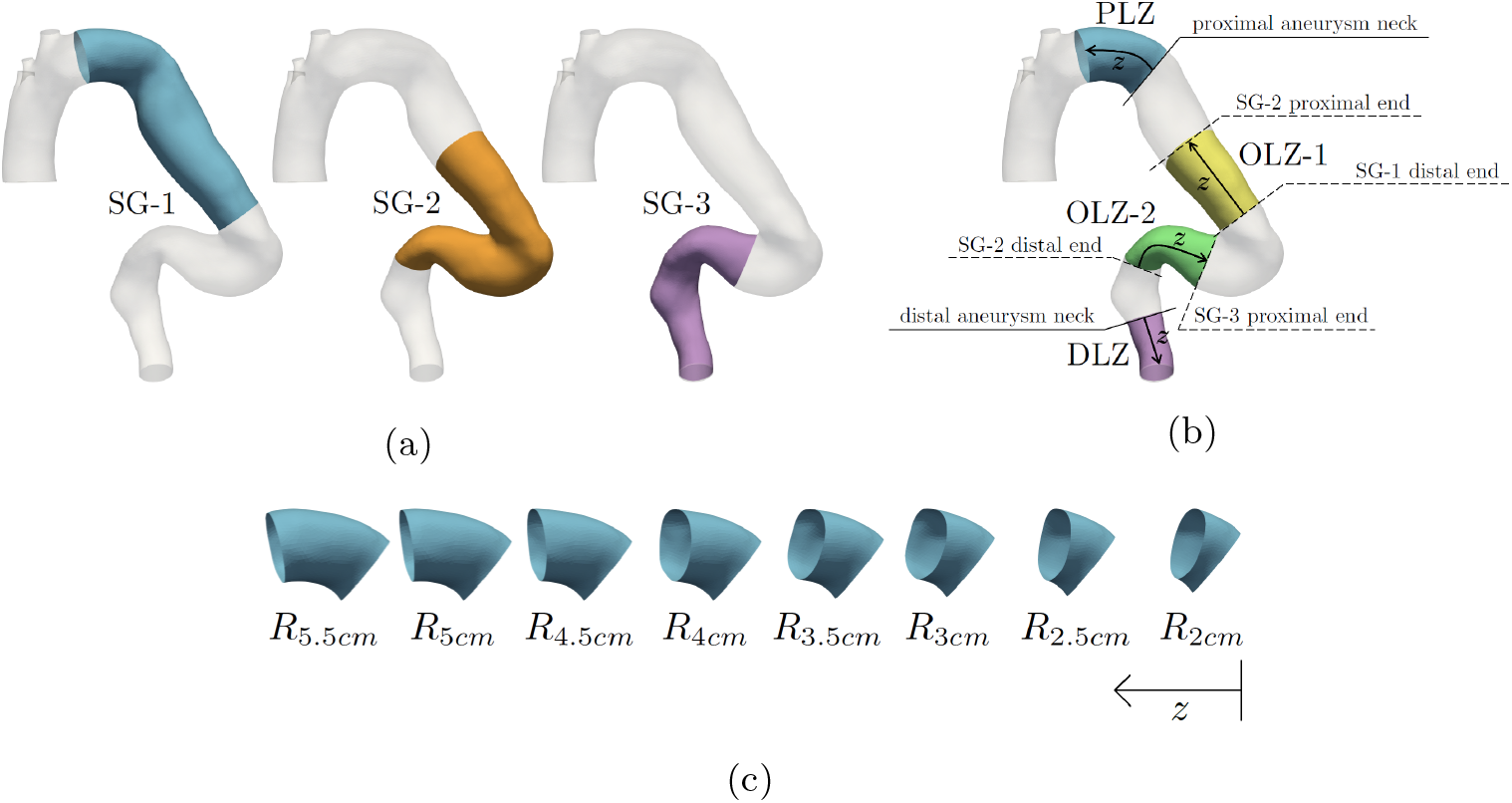
Workflow for the computation of the risk factor *R* in PT-67 (chosen here as representative). (a) Three stent-grafts are present: SG-1, SG-2 and SG-3. (b) Landing zones considered for the analysis: proximal landing zone PLZ, distal landing zone DLZ, overlap landing zones OLZ-1 and OLZ-2. Arrows indicate the direction used to progressively extend each landing zone. (c) Incremental segments of PLZ used for the computation of the risk factor *R*.

Figure 4 displays the entire workflow followed to compute the risk factor *R*.In order to assign specific risk levels to given *R*, we split the patient cohort into: a *Calibration set*, formed by 4 patients with no ELs (PT-09, PT-22, PT-23 and PT-78) and 4 with ELs (PT-10, PT-21, PT-50, PT-51), and a *Validation set*, composed by 1 patient without ELs (PT-40) and 1 with EL (PT-67) (see Table 1 and Figure 1a).The first sub-group is used to associate specific ranges of *R* values to three different levels of risk: *low, moderate* and *high*. This classification is then applied blindly to the second sub-group of patients in order to validate its efficacy through the post-TEVAR follow-up outcomes.

## 3 Results

In Section 3.1 we report the results of the calibration of the risk factor values; in Section 3.2 we present the results of the validation against the post-TEVAR follow-up outcomes (see Section 2.4).

### 3.1 Results for the calibration set

Table 3 reports the values of the risk factor *R*, introduced in Section 2.4, computed for the Proximal Landing Zone (PLZ), Distal Landing Zone (DLZ) and Overlap Landing Zone (OLZ) of all patients of the calibration set (see Table 1 and Figure 1a). The table show that all LZs affected by Endoleaks (ELs) exhibits *R* values greater than 0.9, with the exception of OLZ of PT-10, which shows a slightly lower value of 0.71. Conversely, nearly all LZs without complications have values below 0.20, except for PLZ of PT-09 (*R* = 0.62) and PT-22 (*R* = 0.39), and one of the two OLZ of PT-51 (*R* = 1.00). Given such considerations, we associate a low risk of EL formation for LZs with *R ≤* 0.33, a moderate risk for those with 0.33 *< R ≤* 0.67, and a high risk for LZs with *R >* 0.67.

**Table 3:** *R* values in the proximal (PLZ), distal (DLZ) and overlap (OLZ) landing zones for all the patients of the calibration set. The post-TEVAR follow-up outcomes (i.e., endoleak presence (✓) or absence (✗)) are reported in the right semi-column under PLZ, DLZ, OLZ-1 and OLZ-2. Values not satisfying the calibration procedure are highlighted in red.

| ID | PLZ |  | DLZ |  | OLZ-1 |  | OLZ-2 |  |
| --- | --- | --- | --- | --- | --- | --- | --- | --- |
| | $R$ | EL | $R$ | EL | $R$ | EL | $R$ | EL |
| PT-09 | 0.62 | ✗ | 0.00 | ✗ | 0.00 | ✗ | - | - |
| PT-10 | 0.00 | ✗ | 1.00 | ✓ | 0.71 | ✓ | - | - |
| PT-21 | 0.94 | ✓ | 0.00 | ✗ | 0.11 | ✗ | - | - |
| PT-22 | 0.39 | ✗ | 0.04 | ✗ | 0.14 | ✗ | - | - |
| PT-23 | 0.02 | ✗ | 0.18 | ✗ | - | - | - | - |
| PT-50 | 0.15 | ✗ | 0.00 | ✗ | 0.00 | ✗ | 1.00 | ✓ |
| PT-51 | 1.00 | ✓ | 0.08 | ✗ | 0.96 | ✓ | 1.00 | ✗ |
| PT-78 | 0.00 | ✗ | 0.00 | ✗ | 0.01 | ✗ | - | - |

Figure 5 displays the variation of *R* with the length of each incremental segment (see Section 2.4) of PLZ and DLZ considered for the patients of the calibration set, resulting in a total of 16 curves (one PLZ plus one DLZ for each patient). According to the figure, while extending the length of the incremental segments of DLZ appears to reduce *R*, extending those of PLZ does not always yield the same outcome; in some curves (i.e., PLZ and DLZ of PT-22, PT-23 and PT-51), it is actually associated with a greater risk factor. In 6 out of 16 curves presented (i.e., DLZ of PT-09 and PT-50, PLZ of PT-21 and PT-78, and PLZ and DLZ of PT-10), a 2-cm incremental segment is associated with high risk of type I EL formation; in 3 curves (i.e., PLZ of PT-09 and PT-23, and DLZ of PT-21) it is associated with a moderate risk, while in the remaining 7 curves (i.e., PLZ of PT-22, PT-50 and PT-51, and DLZ of PT-22, PT-23, PT-51 and PT-78) it is associated with low risk.

**Fig. 5:**
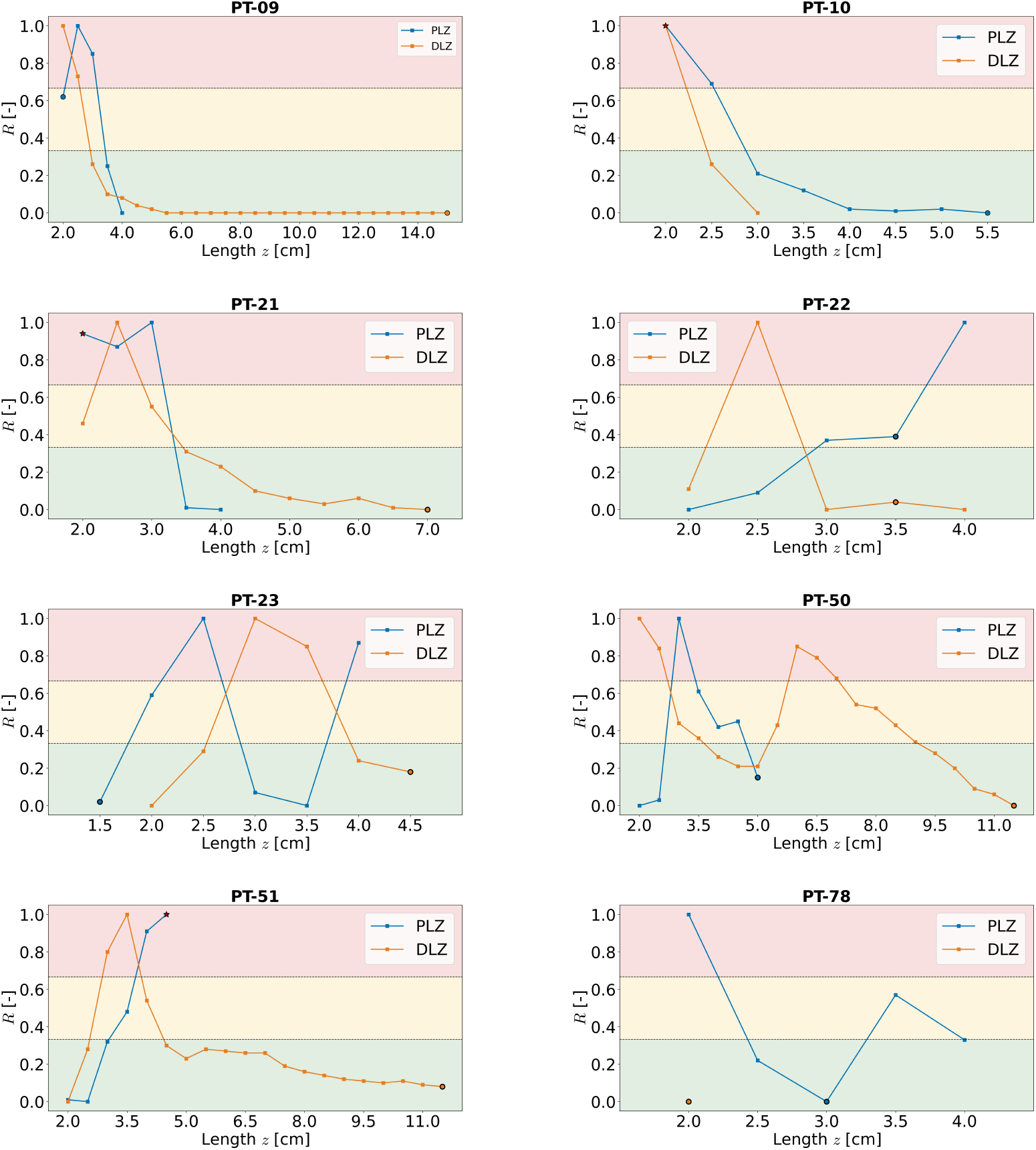
*R* values in all the incremental segments of the proximal (PLZ) and distal (DLZ) landing zones considered for the calibration set as a function of the local coordinate *z*. Circular markers indicate the actual landing zone selected by the clinicians during the intervention. Red star-shaped markers identify the actual landing zone where either a type IA (proximal) or a type IB (distal) endoleak occurred. Horizontal dashed lines delineate the three risk levels: low (green), moderate (yellow) and high (red).

Figure 6 shows the variation of *R* with the length of each incremental segment of all OLZs considered for the patients of the calibration set, resulting in a total of 9 curves (see Table 1). According to the figure, in 5 out of 9 curves presented (i.e., OLZ of PT-09, PT-21, PT-22 and PT-78, and OLZ-1 of PT-50), increasing the length of the incremental segments of OLZ leads to a progressive reduction in *R*, reaching values associated with low risk of type III EL formation. On the contrary, in the remaining 4 curves (i.e., OLZ of PT-10, OLZ-2 of PT-50, and OLZ-1 and OLZ-2 of PT-51), the progressive extension of OLZ results in an increase in *R*, reaching values associated with high-risk condition. Moreover, in 6 curves (i.e., OLZ of PT-10, PT-21, PT-22 and PT-78, OLZ-2 of PT-50, and OLZ-1 of PT-51), a 2-cm incremental segment is associated with a low risk of EL occurrence, in 2 curves (i.e., OLZ of PT-09, and OLZ-2 of PT-51) with moderate risk, and in the remaining curve (i.e., OLZ-1 of PT-50) with high risk.

**Fig. 6:**
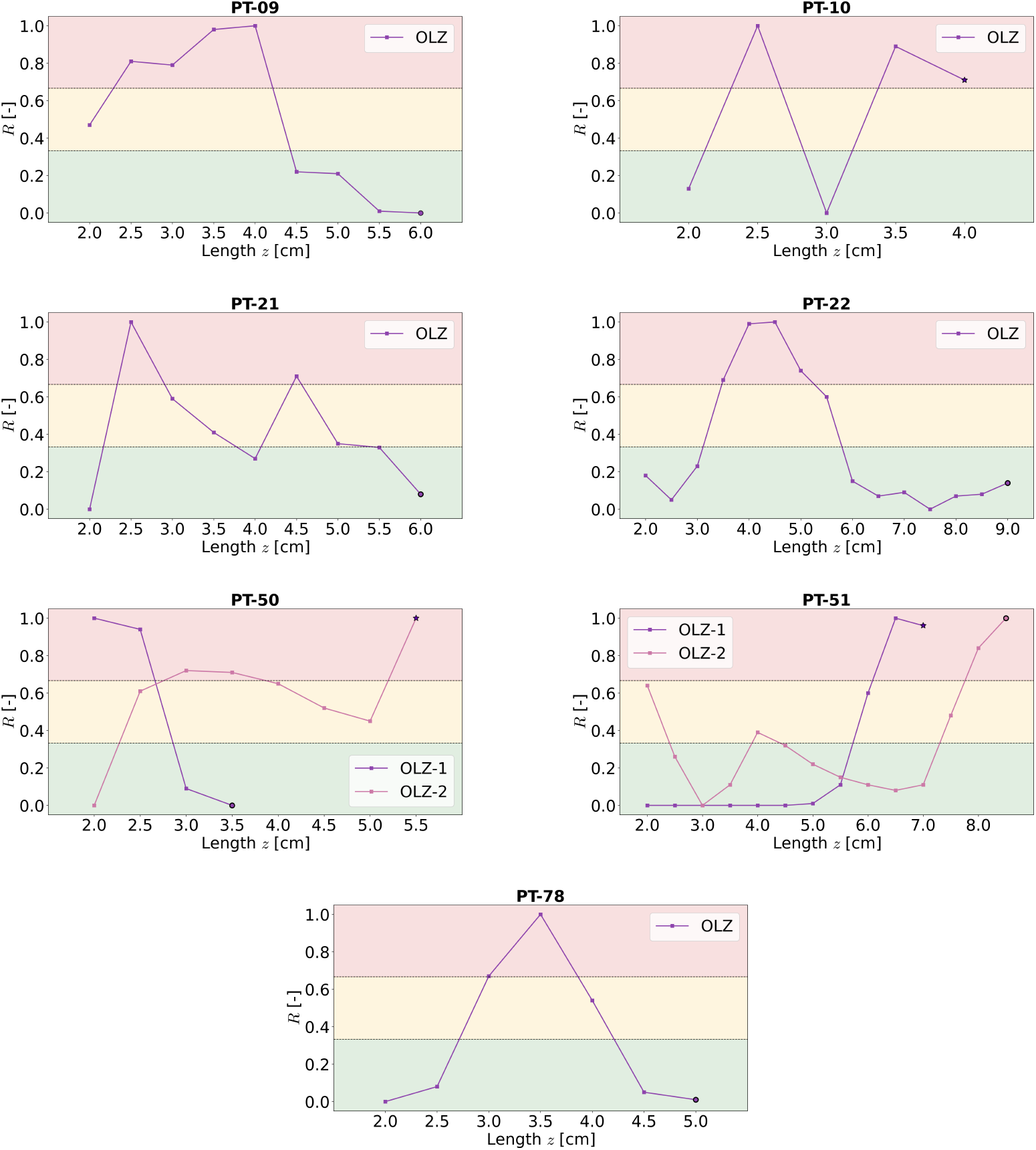
*R* values in all the incremental segments of the overlap (OLZ) landing zones considered for the calibration set as a function of the local coordinate *z*. Circular markers indicate the actual landing zone selected by the clinicians during the intervention. Purple star-shaped markers identify the actual landing zone where a type III endoleak occurred. Horizontal dashed lines delineate the three risk levels: low (green), moderate (yellow) and high (red).

### 3.2 Results for the validation set

Table 4 reports the values of the calibrated risk factor *R* computed for PLZ, DLZ and OLZ of patients not used for the calibration procedure, i.e. PT-40 and PT-67 belonging to the validation set. According to the table, *R* is able to correctly predict the occurrence of type Ib EL in PT-67, since it is equals to 1.00 in DLZ. Moreover, *R* prescribes a low-risk condition in PLZ and two OLZs, in good accordance with the follow-up condition of PT-67. On the other hand, PT-40 had no complications, and *R* accordingly predicts a low risk of EL formation in PLZ, DLZ and OLZ.

**Table 4:**
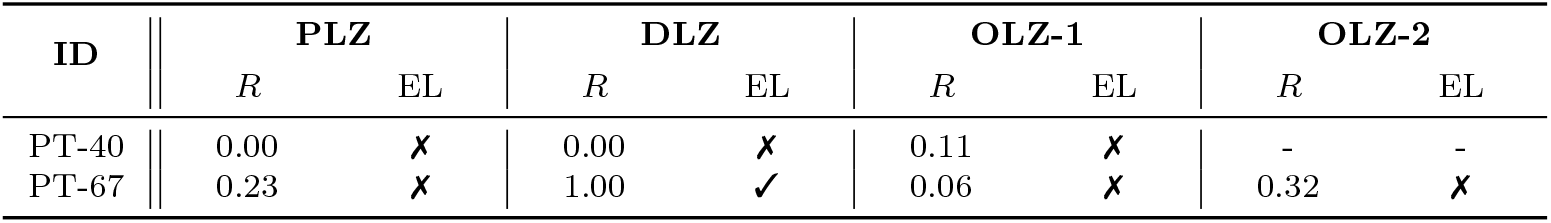
*R* values in the proximal (PLZ), distal (DLZ) and overlap (OLZ) landing zones for all the patients in the validation set. The post-TEVAR follow-up outcomes (i.e., endoleak presence (✓) or absence (✗)) are reported in the right semi-column under PLZ, DLZ, OLZ-1 and OLZ-2.

| ID | PLZ |  | DLZ |  | OLZ-1 |  | OLZ-2 |  |
| --- | --- | --- | --- | --- | --- | --- | --- | --- |
| | $R$ | EL | $R$ | EL | $R$ | EL | $R$ | EL |
| PT-40 | 0.00 | ✗ | 0.00 | ✗ | 0.11 | ✗ | - | - |
| PT-67 | 0.23 | ✗ | 1.00 | ✓ | 0.06 | ✗ | 0.32 | ✗ |

Figure 7 displays the variation of *R* with the length of each incremental segment of PLZ, DLZ and OLZs considered for the patients of the validation set, resulting in a total of 7 curves (see Table 1a). In all the curves presented, increasing the length of the incremental segments leads to a decrease in *R*, and consequently with a lower risk of both type I and III EL formation. Moreover, for all OLZs analysed, a 2-cm incremental segment is associated with low risk of type III EL. Conversely, for all the distal regions, a 2-cm incremental segment corresponds to a high-risk condition, whereas for PLZ, it corresponds to high risk in PT-67, and moderate risk in PT-40.

**Fig. 7:**
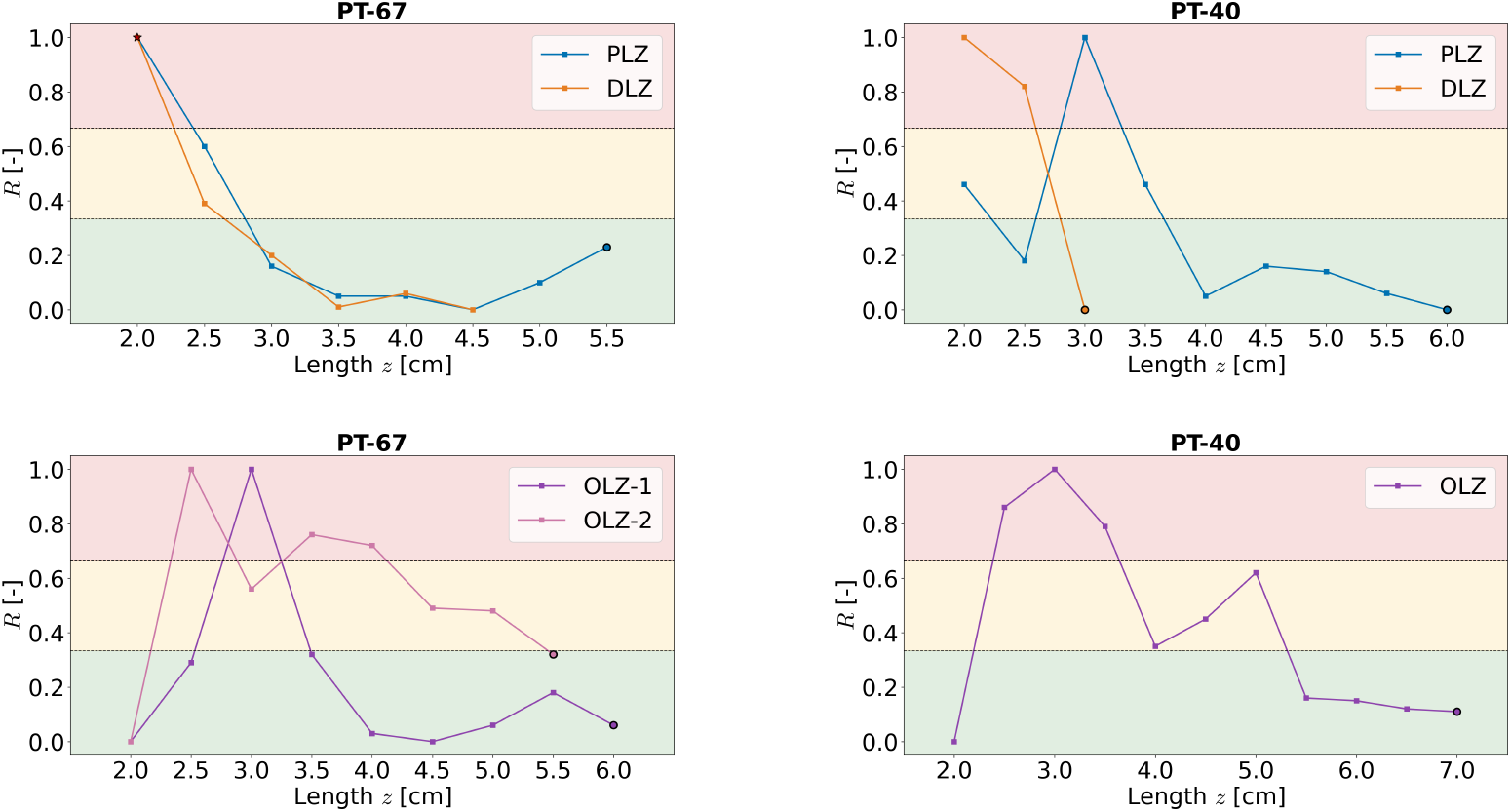
*R* values in all the incremental segments of all the landing zones considered for the validation set as a function of the local coordinate *z*. Top row: proximal (PLZ) and distal (DLZ) landing zones. Bottom row: overlap (OLZ) landing zones. Circular markers indicate the actual landing zone selected by the clinicians during the intervention. Red star-shaped markers identify the actual landing zone where either a type IA (proximal) or a type IB (distal) endoleak occurred. Purple star-shaped markers identify the actual landing zone where a type III endoleak occurred. Horizontal dashed lines delineate the three risk levels: low (green), moderate (yellow) and high (red).

## 4 Discussion

This study proposes a risk factor *R* able to predict post-TEVAR type I and III ELs occurrence only on the basis of the patient-specific pre-TEVAR hemodynamics simulated within an FSI framework.

The results obtained in the calibration set (see Table 3) indicate that the risk factor *R* is able to well capture the occurrence of adverse events, showing very high values (usually greater than 0.90) in cases with ELs, and very low values (less than 0.20) in cases without complications. There are only two exceptions: (i) PLZ of PT-09 (*R* = 0.62) and PT-22 (*R* = 0.39), which exhibit slightly elevated values despite the absence of EL, and (ii) one of the two OLZs of PT-51 (*R* = 1.00), which is also not subjected to complications.

The first exception is likely related to the fact that the zone selected for the proximal landing is located in correspondence of the Aortic Arch (AA) (i.e., Zone 3, see Table 1), which is a region with great curvature, known to be associated with a higher risk of TEVAR-related complications [28]. In this regard, it is worth noting that PT-09, who presents a type III AA (i.e., the most sharply curved), exhibits a higher risk factor compared to PT-22, who has instead a type I AA.

The second exception can be explained by the specific configuration adopted in PT-51. In this case, a main endoprosthesis was implanted with two fenestrations to accommodate two modular stent-grafts, one for the BCA and one for the LCCA. Both modular components overlap with the main device near its PLZ (i.e., Zone 0, see Table 1). Notably, the latter is affected by a type IA EL, while the BCA modular part presents a type III EL. This indicates the presence of particularly adverse hemodynamic conditions near the main stent-graft PLZ. Therefore, since the LCCA modular part overlaps the main device in the same region, it is reasonable to assume that it may be exposed to similar hostile hemodynamics. This is reflected by the high *R* value observed, despite the absence of complications at the time of the patient last follow-up.

Based on the risk factor values obtained in the calibration set, we associate low risk of type I and III EL formation for LZ with *R ≤*0.33, moderate risk for those with 0.33 *< R ≤*0.67, and high risk for LZs with *R >* 0.67. Therefore, overall, the only case mispredicted by the risk factor is OLZ-2 in patient PT-51, since PLZ of patients PT-09 and PT-22 lies within the moderate-risk level, where endoleak occurrence is not certain (see Table 3).

When the risk-level classification is applied blindly to the validation set (see Table 4), a single high-risk *R* value is encountered in DLZ of PT-67, where a type Ib EL actually occured. In all the other regions, for both PT-67 and PT-40, the risk factor predicts low risk of EL formation. Therefore, these findings seem to support the validity of the proposed risk factor as a tool for identifying, before the intervention, cases at higher risk of EL formation.

In general, according to the results (see Figure 5 and Figure 7), it appears that extending DLZ to move as far as possible from the aneurysmal sac (which is a region characterized by elevated drag forces [35]) always tends to reduce the risk of type Ib EL. Thus, in PT-10, the type Ib EL might have been avoided by considering a longer DLZ, as the actual one may not have been sufficiently long to move away from the aneurysm. Although in 4 out of 10 patients (i.e., PT-09, PT-10, PT-40 and PT-67) the risk decreases monotonically with increasing DLZ length, in the remaining patients, localized peaks are still observed. These localized risk intensifications are probably due to the presence of anatomically challenging regions, such as areas characterized by high tortuosity or pronounced curvature.

On the contrary, selecting a very long PLZ does not necessarily translate into a lower risk of type Ia EL formation (see PT-22, PT-23 and PT-51 in Figure 5). This can be explained by the fact that stent-graft deployment in TEVAR typically requires a proximal landing on the AA (i.e., Zone 0, 1, 2 and 3), which is known to be a hostile region [53]. Indeed, as it advances along the TA towards the heart, the endograft is increasingly subjected to intense pulsatile blood flow generated by the cardiac ejection. Moreover, TA presents a three-dimensional angulation between the AA and DTA, as well as the supraortic vessels arising at short intervals along the arch. All these factors make it difficult to obtain a sufficiently secure PLZ for stent-graft fixation [54]. Therefore, to mitigate the risk of type Ia EL, it is necessary to ensure that, as the PLZ is progressively extended in order to move far from the aneurysm, any potential increase in drag forces, due to the inclusion of hemodynamically hostile regions, is fully compensated by the corresponding increase in contact area between the stent-graft and the aortic wall.

A similar conclusion can be drawn for OLZ (see Figure 6 and Figure 7): a longer LZ is not necessarily associated with a lower risk of type III EL (as observed in PT-10, PT-50 and PT-51). This is probably due to the fact that the overlap between two stent-grafts is generally performed within the aneurysmal sac: a region characterized by intense drag forces and particularly disturbed hemodynamics [35]. Therefore, it is more likely that, when the OLZ length is increased, regions characterized by progressively greater hemodynamic forces are included and, if these forces are not adequately counterbalanced by the larger contact area between the two stent-grafts, the risk of type III EL formation may increase.

This study represents a first attempt to investigate a possible relationship between pre-operative hemodynamics and post-TEVAR complications. The results presented suggest that the pre-TEVAR hemodynamics may provide predictive insights into TEVAR outcomes. In particular, the risk factor proposed in this work seems to be capable of identifying all the EL cases analysed, as well as conditions known to pose a risk for procedural complications. This risk factor could be useful not only for predicting the risk of EL formation, associated with a specific LZ already selected by the surgeons, but also for helping clinicians identify, on a patient-specific basis, the safest region for the stent-graft sealing, thereby potentially contributing to improve TEVAR outcomes.

This study presents some limitations:

- the pressure-wave employed as inlet BC in the FSI simulations is taken from literature, and it is not patient-specific. Prescribing patient-specific inlet condition can be achived by exploiting either Echo Doppler cardiac output or catheter-based pressure measurements. This could not be done in the present work due to the retrospective nature of the study;
- the aortic wall is modelled as a linear elastic material, even though the human aorta exhibits hyperelastic and viscoelastic behaviour. Nevertheless, as the proposed risk factor is based exclusively on blood forces, we consider this modeling assumption as reasonable;
- the proposed risk factor is based only on the hemodynamics, but it would be of interest to consider also structural characteristics, such as von Mises Stresses;
- the number of patients analysed is probably too small to draw statistically significant conclusions. However, it is sufficient for a first exploratory attempt to study the relationship between pre-operative hemodynamics and TEVAR outcomes. In the future, we intend to expand the patient cohort and conduct a prospective study, in which the optimal stent-graft LZs are selected based on this analysis.

## Author contributions

F.D.: conceptualization of the work, development of the methodology, code development, images reconstruction, numerical simulations, post-processing of the results, interpretation of the results, writing of the original draft, editing of the final version of the manuscript. Si.T.: images reconstruction, numerical simulations, post-processing of the results. M.D.: conceptualization of the work, acquisition of the clinical data, interpretation of the results, revision of the manuscript. D.B.: conceptualization of the work, acquisition of the clinical data, interpretation of the results, revision of the manuscript. Sa.T.: conceptualization of the work, acquisition of the clinical data, interpretation of the results, revision of the manuscript. C.V.: conceptualization of the work, development of the methodology, interpretation of the results, revision of the manuscript, supervision, funding acquisition. F.M.: conceptualization of the work, development of the methodology, interpretation of the results, revision of the manuscript, supervision, funding acquisition.

## Acknowledgements

F.D. and C.V. are members of the INdAM group GNCS ‘Gruppo Nazionale per il Calcolo Scientifico’ (National Group for Scientific Computing). C.V. has been partially supported by (i) the European Union-Next Generation EU, Mission 4, Component 1, CUP: D53D23018770001, under the research project MIUR PRIN22-PNRR n. P20223KSS2, ‘Machine learning for fluid structure interaction in cardiovascular problems: efficient solutions, model reduction, inverse problems’; (ii) the Italian Ministry of Health within the PNC PROGETTO HUB LIFE SCIENCE - DIAGNOSTICA AVANZATA (HLS-DA) ‘INNOVA’, PNCE3-2022-23683266, CUP: D43C22004930001, within the ‘Piano Nazionale Complementare Ecosistema Innovativo della Salute’, Codice univoco investimento: PNCE3-2022-23683266; (iii) Italian Ministry of Health within the project ‘CAL.HUB.RIA’ - CALABRIA HUB PER RICERCA INNOVATIVA ED AVANZATA, Code: T4-AN-09, CUP: F63C22000530001. The authors acknowledge the CINECA award under the ISCRA initiative, for the availability of high-performance computing resources and support.

## Declarations

### Ethical Approval

Informed consent was obtained from all patients.

### Open Access

This article is licensed under a Creative Commons Attribution 4.0 International License, which permits use, sharing, adaptation, distribution and reproduction in any medium or format, as long as you give appropriate credit to the original author(s) and the source, provide a link to the Creative Commons licence, and indicate if changes were made. The images or other third party material in this article are included in the article’s Creative Commons licence, unless indicated otherwise in a credit line to the material. If material is not included in the article’s Creative Commons licence and your intended use is not permitted by statutory regulation or exceeds the permitted use, you will need to obtain permission directly from the copyright holder. To view a copy of this licence, visit http://creativecommons.org/licenses/by/4.0/.

## Footnotes

1 the average of the signal over the seven heartbeats considered

